# Metric Mirages in Cell Embeddings

**DOI:** 10.1101/2024.04.02.587824

**Authors:** Hanchen Wang, Jure Leskovec, Aviv Regev

## Abstract

Although biological studies increasingly rely on embeddings of single cell profiles, the quality of these embeddings can be challenging to assess. Such evaluations are especially important for avoiding misleading biological interpretations, assessing the accuracy of integration methods, and establishing the zero-shot capabilities of foundational models. Here, we posit that current evaluation metrics can be highly misleading. We show this by training a three-layer perceptron, Islander , which outperforms all 11 leading embedding methods on a diverse set of cell atlases, but in fact distorts biological structures, limiting its utility for biological discovery. We then present a metric, scGraph, to flag such distortions. Our work should help learn more robust and reliable cell embeddings.

---

Embeddings of single cell profiles are now routinely employed as a research tool in biological investigation, to characterize cell types and states, their changes over time, and their distinction between conditions, including diseases, organs, or drug treatments (1, 2). With a dramatic growth in single cell data, including the Human Cell Atlas (3), multiple efforts have focused on universal embeddings for diverse single cell data (4–9). Given their broad utility, it is crucial to scrutinize the embeddings’ quality to evaluate the performance of the underlying integration method (10) and zero-shot capabilities of the resulting foundation model (11, 12).

A critical aspect in deriving helpful cell embeddings is the correction of non-biological batch effects that stem from technical variations, such as sample handling and sequencing protocols. These variations can mask biological signals and lead to misleading interpretations. Integration methods thus aim to mitigate batch-specific discrepancies while preserving essential biological information. The effectiveness of these integrated cell embeddings is typically assessed through two lenses: how well the cells from various batches mix together; how closely cells of the same type group together.

Here, we identified an overlooked challenge in the evaluation metrics used to assess embeddings. To demonstrate the limitations of current gold standard metrics for cell profile embeddings (10), we first developed Islander (Fig. 1a), a model that scores best on established metrics, but generates biologically problematic embeddings. Islander is a three-layer perceptron, directly trained on cell type annotations with mixup augmentations (13). We tested Islander across a diverse set of 11 different human tissue cell atlases (brain (14), breast (15), eye (16), fetal gut (17), heart (18), fetal lung (19), pancreas (10), skin (20)), which cover different strengths of batch effects and diverse biological systems, overall comprising more than 3.5 million cells from 10 human organ systems. For each atlas, we trained a Islander model and then compared it with another 12 embedding baselines: three dimension reduction methods (PCA, UMAP, TSNE) (21), eight batch integration methods (Harmony (22), Scanorama (23), BBKNN (24), fastMNN (25), scVI (26), scANVI (27), scGen (28), scPoli (29)), and one foundation model (Geneformer (5)) (**Methods**). In addition, for each atlas, we compared to the performance of the original authors’ integration, if available.

**Figure 1:**
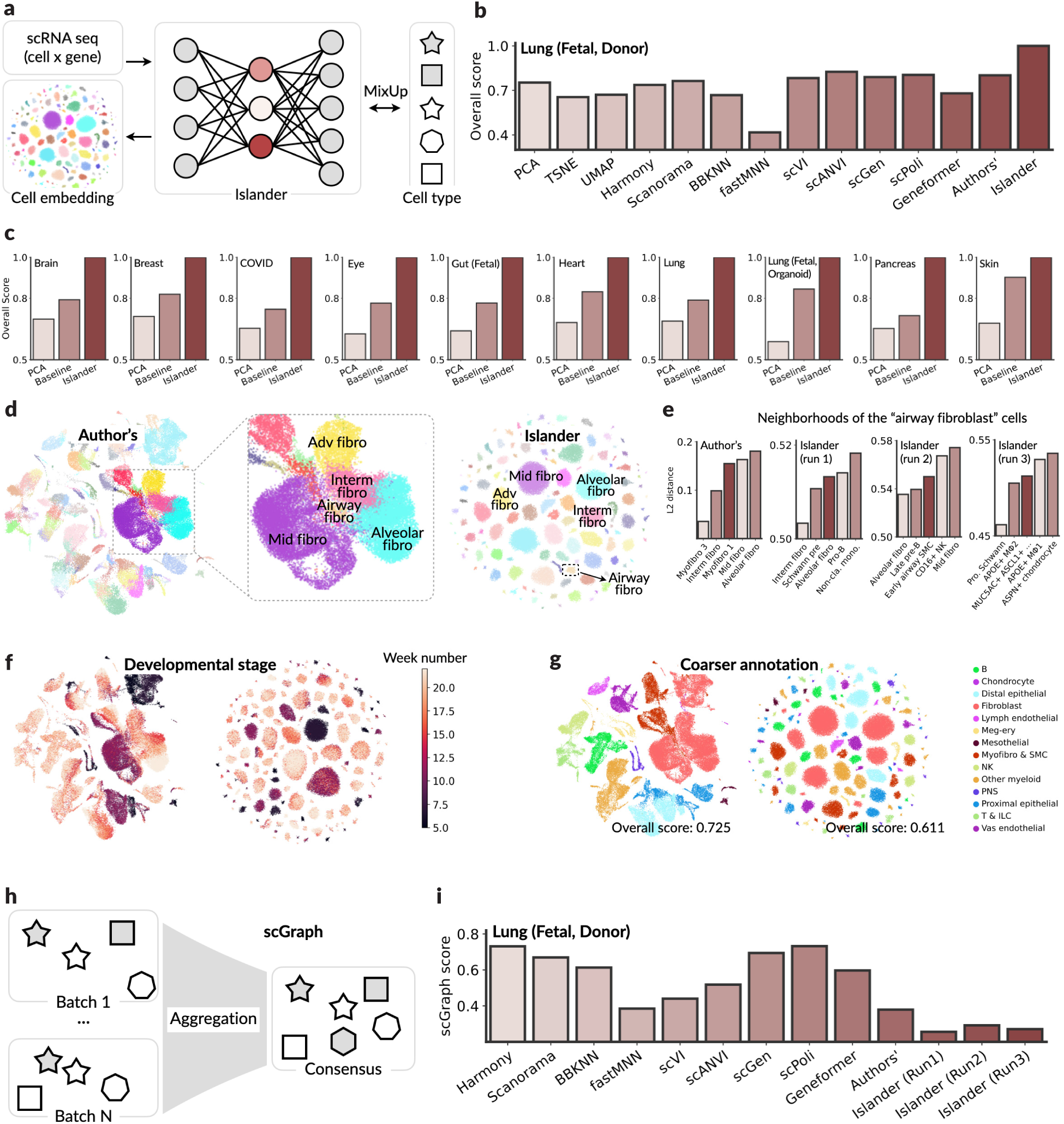
Drifting Cell Islands highlight limitation of current metrics. **a,** Islander overview. **b, c,** Evaluation of cell embeddings. Normalized overall score over 12 metrics (y axis, **Methods**) for each method (x axis) assessed on using the Fetal Lung Cell Atlas (b) or another 10 cell atlases (c). “Baseline“: best baseline results (**Methods**). **d,** Fetal Lung Cell Atlas embedding space. Single cell profiles (dot; color-coded by cell type annotation) from the Fetal Lung Cell Atlas embedded by the author’s integration method (left, inset zoom in middle) or Islander (right). Annotations: fibroblast cell subsets. **e,** “Airway fibroblast” cell neighborhood changes across Islander runs. Normalized Euclidean distance (y axis) between the centroids of airway fibroblasts and its five nearest neighbor clusters (x axis) in the 50-dimensional (of author’s integration) and 16-dimensional (of Islander ’s) embedding space. **g,** Cell islands distort developmental stage structure and cell cell relationships. Cell embeddings as in (d), colored by developmental week (f, color bar) or coarser cell type annotations (g). **h, i.** scGraph, a metric using learned cell similarity graphs to evaluate cell embeddings. h. Method overview. i. scGraph score (y axis, higher is better) for each method (x axis) assessed using the Fetal Lung Cell Atlas.

Across all datasets, Islander consistently outperformed all baseline strategies across all 12 metrics (10) (Fig. 1b,c, Extended Data Tables 3-13). This is largely due to the principles underlying the evaluation metrics (10), which focus on assessing the efficiency of cell embeddings in terms of the coherence of cell clustering structures with cell type labels and the blending of batches within clusters. When Islander explicitly aligns these cell embeddings with cell type annotations, it forms well-separated cell ‘islands’ (Fig. 1d, right), with each island comprising cells annotated as the same type. This alignment significantly boosts the biological variance conservation metrics, leading to top-tier overall performance (Extended Data Tables 3-13).

However, such structure is driven by (and complies well) with the most granular annotation level at the cost of ignoring any higher level relationships and distorting biological structures, potentially obstructing downstream analyses and future discoveries (and would thus not be an advisable for an actual integration method). In particular, when annotated cell subsets follow a continuum, as is the case for fibroblasts, Islander separates its constituent parts (Fig. 1d). In the developing human lung, the original analysis (19) identified multiple sub-types of fibroblasts, each distinguished by different marker genes and spatial locations. While the original embedding preserves a continuum between these fibroblasts (Fig. 1d, left), they are fully separated by Islander (Fig.1d, right). Similarly, the Islander embedding disrupted the developmental continuum, clearly observed in the original study (Fig. 1f, left), but obscured by Islander (Fig. 1f, right).

Moreover, the “cell islands” drifted in different ways across distinct runs, especially for smaller cell subsets. For example, in three separate runs with overall similar scores, the composition of the neighborhood of airway fibroblast cells varied substantially, involving as many as 14 distinct cell types within the five nearest neighbors (Fig. 1e, Extended Data Fig. 1, Extended Data Table 9). Thus, aside from cluster identity, the embedding may be largely arbitrary in all other relationships, and this arbitrariness would carry into downstream analysis or the biologist’s interpretation.

Prompted by these limitations of the quality evaluation criteria, we reasoned that focusing solely on the most granular cell relationships in evaluation can pose a substantial limitations, whereas preserving relationships between broader cell types (coarser annotations) is an important additional criterion, and may also be more robust to noise. Indeed, when evaluating the same set of embeddings using broader cell type annotations provided by the authors, Islander now achieved an overall score of 0.523, inferior to PCA (0.557) or the top-performing scVI (26) (0.701) (Extended Data Table 14).

Because hierarchical Cell Ontology annotations are often unavailable (6), we next developed scGraph, as a new framework for quality assessment (**Methods**). For each set of cell embeddings, we define an affinity graph to elucidate the similarities between various cell types. scGraph then compares each affinity graph to a consensus graph, derived by aggregating individual graphs from different batches, based on raw reads or PCA loadings. This metrics efficiently highlights the inherent biological structures, emphasizing cell type similarities, while reducing the impact of technical variations across batches. Notably , scGraph does not require that any single batch have cells of all types (Fig. 1h), thus fitting the constraints of real datasets in the domain.

Evaluation by scGraph revealed varied performance across embeddings. Here, Islander had lower scores (Fig. 1i, Extended Data Table 15), while Harmony and scPoli excelled in capturing the complex relationships within functional cellular clusters. Indeed,Islander was the lowest scoring across seven of 11 atlases, underscoring scGraph’s ability to detect the “drifting cell islands” artifact. Interestingly, scGraph favored higher-dimensional embeddings, like PCA over a PCA-derived UMAP. Note that scGraph’s premise that profiles of functionally-similar cells would be proximate, may not always hold true.

In conclusion, we demonstrated the limitation of current quality metrics by introducing Islander , a three-layer perceptron, an integration approach that outperforms all major methods across diverse cell atlases, but at the cost of “island-like” distortions in the biological structures in cell embedding spaces. To address the inherent limitations of current evaluation metrics, we propose a new approach, scGraph, to helps assess how well the results of an integration approach preserve cell cell relationships at multiple granularities. Our work also highlights the importance of incorporating weaker supervision, as was recently illustrated in approaches where an encoder is regularized with an additional reconstruction loss (6), or using large-scale unsupervised pre-training (*e.g.*, Universal Cell Embedding (UCE) (7)). These advancements underline the significance of methodological choices in computational biology and offer guiding principles for future research.

## Methods

### Datasets and pre-processing

Raw sequencing data were downloaded from the respective data providers as of October 1, 2023; details on the datasets and their sources are provided in Extended Data Table 1. The analysis encompasses 11 cell atlases, totaling 3,510,450 cell profiles. A uniform pre-processing protocol was applied across these datasets. Specifically, cell profiles with fewer than 1,000 reads or less than 500 detected genes were filtered out, and genes present in fewer than five cells were also excluded. Normalization was performed using Scanpy (30), scaling each cell’s read counts to a total of 10,000 and subsequently applying a log1p transformation.

### Baselines

Eleven baseline methods were used for comparison: three dimensionality reduction baselines: PCA, t-SNE (31), and UMAP (32); eight integration methods: Harmony (22), BBKNN (24), Scanorama (23), fastMNN (25), scVI (26), scANVI (27), scGen (28), and scPoli (29); and one pretrained foundation model: Geneformer (5), for zero-shot embedding extraction. For dimensionality reduction methods, the log1p transformed raw counts from gene-by-cell matrices were provided as input. For each integration method, default hyperparameter settings recommended by the original authors were used. For Geneformer, the largest pre-trained model weights provided by the authors (33) were used. While scANVI, scGen, and scPoli utilize cell type as parts of their computational pipelines, other integration methods do not require such information. The top 1,000 genes were identified as highly variable gene sets.

### Assessment metrics

Cell embeddings was assessed using metric as described in previous study (10), and implemented in “scib-metrics” (34). The following evaluation metrics were used (abbreviations noted are used in Extended Data Tables 3-13) . ’I-label’ for isolated labels, ’L-NMI’ for Leiden normalized mutual information, ’L-ARI’ for Leiden Averaged Rand Index, ’K-NMI’/’K-ARI’ for K-means NMI/ARI, ’S-label/batch’ for silhouette label/batch, ’c/i-LISI’ for batch-mixing (iLISI) and cell-type separation (cLISI), ’G-Con’ for graph connectivity, and ’PCR’ for principal component regression. Consistent with previous studies, selection of highly variable genes enhanced the performance of data integration methods.

### Islander design

Islander is as a three-layer perceptron with two hidden layers of sizes 128 and 16, respectively, and an output layer matching the total number of cell types as annotated. The first hidden layer incorporates ReLU activation and batch normalization, while cell embeddings are derived from the second hidden layer. The output layer employs a softmax normalization function. In extended experiments, a decoder module was added mirroring the original MLP structure, with output dimensions of 16, 128, and the total number of genes, respectively. Each layer in this extended setup uses ReLU activation and batch normalization, except for the final linear layer.

### Training setup

The model was trained in a manner aligned with scvi-tools (35), with mini-batches of 256 randomly sampled cells from all batches, along with their cell type annotations. Islander was trained using cross-entropy loss with mixup (13) augmentations (default setting). The Adam optimizer was used with an initial learning rate of 0.001, over 10 epochs, and a cosine annealing scheduler for learning rate decay. All cells were utilized for training to maximize overfitting.

### Neighborhood calculation

Neighborhoods of each cell type were identified by the Euclidean distance between the centroids of cell profiles of each type in the embedding space. To mitigate the effects of batch variation and measurement noise, a trimming strategy was applied. The outermost 20% of data, treated as outliers, were excluded before calculating the centroid coordinates. This ensures a more accurate representation of cell type proximity by focusing on the most representative data points.

### scGraph

scGraph quantifies the similarity between two graphs that each represent the closeness between cell types. In these graphs, each entry (*x*, *y*) signifies the proximity of cell type *x* to cell type *y*. The first graph is derived from the provided embeddings, while the second, serving as a reference, is based on raw counts or PCA loadings from each batch. For the reference, proximity graphs are initially computed from each batch using normalized Euclidean distances between centroids of the cell type profiles. These batch-specific graphs are then amalgamated into a single consensus graph through averaging. The similarity of neighborhoods for each cell type is assessed using Pearson’s rank correlation. The final score, reflecting the overall similarity and ranging from -1 to 1 (with higher values indicating greater similarity), is the average across all cell types. The goal is to align the neighborhood graphs from the embeddings with the reference graph derived from the data, indicating that cells with similar profiles are appropriately clustered in the embedding space.

### Code availability

The implementation code for the Islander , as well as the tutorial notebooks sufficient to reproduce the results presented in this manuscript, can be accessed via https://github.com/Genentech/Islander. For scIB evaluation pipelines, we use the implementations by Gayso et al from https://github.com/yoseflab/scib-metrics.

## Acknowledgment

We thank Romain Lopez, Peng He, Leander Dony, Sara-Jane Dunn, Gocken Eraslan, Adam Gayoso, Graham Heimberg, Kexin Huang, John Marioni, Dana Pe’er, Yusuf Roohani, Yanay Rosen, Andrew Whitehead, and Jiaqi Zhang for invaluable insights, along with other members of the Leskovec and Regev labs and colleagues at the Human Cell Atlas, Chan Zuckerberg Initiative, and Google Deep-Mind for constructive discussions.

## Supplementary

### 1. Data availability

**Extended Data Table 1:** Statistics of cell atlases. “# Class” represents the total number of cell types.

| Atlas | # Gene | # Cell | # Class | # Batch | Reference |
| --- | --- | --- | --- | --- | --- |
| Brain | 59,357 | 888,263 | 11 | 4 | Paper (14), Data source (36) |
| Breast | 33,234 | 703,512 | 39 | 126 | Paper (15), Data source (37) |
| COVID | 33,537 | 559,517 | 31 | 10 | Paper (38), Data source (39) |
| Eye | 36,484 | 51,645 | 11 | 8 | Paper (16), Data source (40) |
| Gut (Fetal) | 26,328 | 62,849 | 21 | 9 | Paper (17), Data source (41) |
| Heart | 33,234 | 486,134 | 27 | 14 | Paper (18), Data source (42) |
| Lung | 28,024 | 584,444 | 53 | 166 | Paper (43), Data source (44) |
| Lung (Fetal, Donor) | 26,354 | 71,752 | 144 | 29 | Paper (19), Data source (45) |
| Lung (Fetal, Organoid) | 24,653 | 70,495 | 28 | 37 | Paper (19), Data source (45) |
| Pancreas | 19,093 | 16,382 | 14 | 9 | Paper (10), Data source (46) |
| Skin | 30,933 | 15,457 | 13 | 5 | Paper (20), Data source (47) |

### 2. Performance

The embedding dimensions of each method is reported in Extended Data Table 2.

**Extended Data Table 2:** Embedding dimensions of each method.

| Method | PCA | TSNE | UMAP | Harmony | Scanorama | BBKNN | fastMNN |
| --- | --- | --- | --- | --- | --- | --- | --- |
| # Dim | 50 | 2 | 2 | 50 | 100 | 2 | 50 |
| Method | scVI | scANVI | scGen | scPoli | Geneformer | Islander |  |
| # Dim | 30 | 30 | 50 | 10 | 512 | 16 |  |

The detailed performance of each cell atlas is shown in Extended Data Tables 3-13. The best aggregated scores are highlighted in **bold**, and the term “Author’s” denotes the author’s integrated embeddings.

**Extended Data Table 3:**
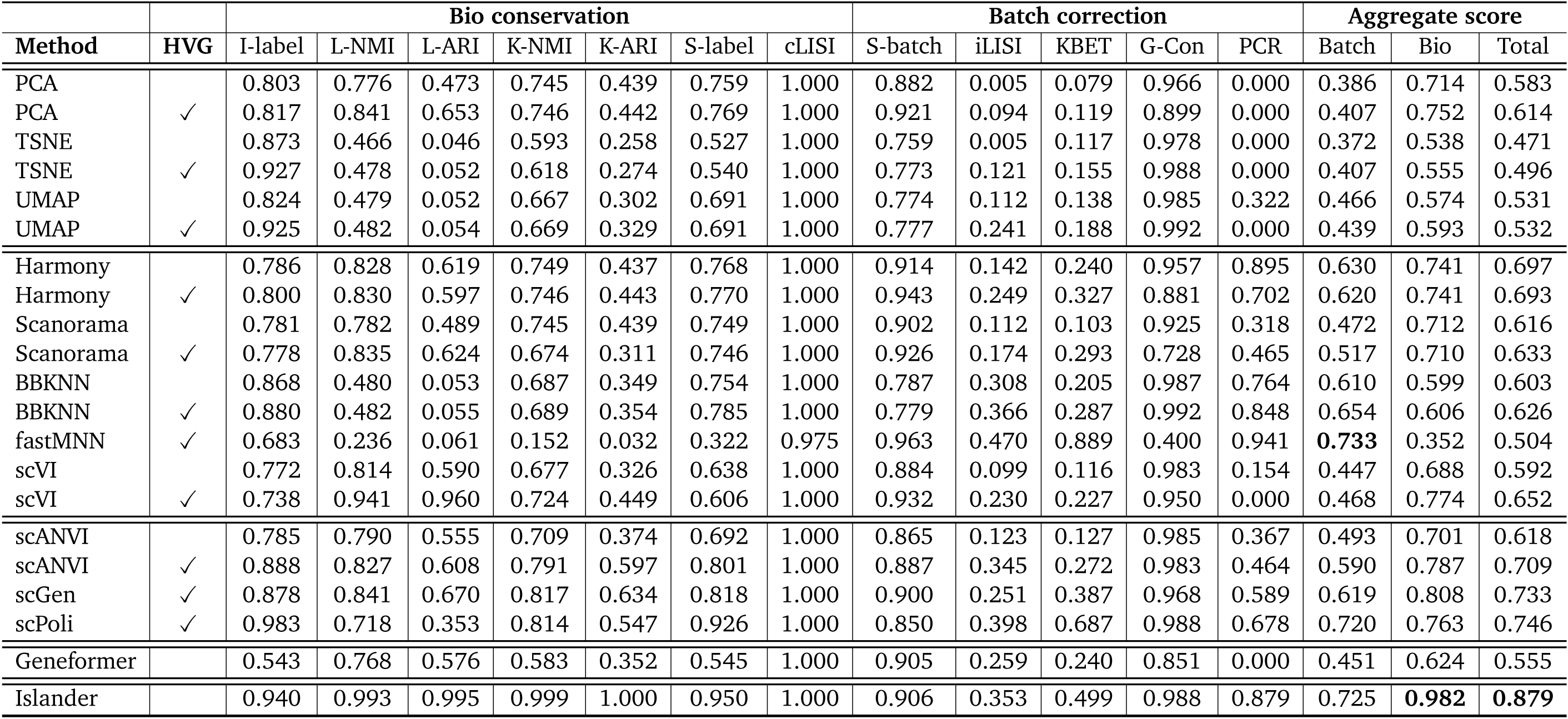
Benchmarking cell embeddings, on Brain Cell Atlas.

**Extended Data Table 4:** Benchmarking cell embeddings, on Breast Cell Atlas.

|  |  | Bio conservation |  |  |  |  |  |  | Batch correction |  |  |  |  | Aggregate score |  |  |
| --- | --- | --- | --- | --- | --- | --- | --- | --- | --- | --- | --- | --- | --- | --- | --- | --- |
| Method | HVG | I-label | L-NMI | L-ARI | K-NMI | K-ARI | S-label | cLISI | S-batch | iLISI | KBET | G-Con | PCR | Batch | Bio | Total |
| PCA |  | 0.656 | 0.776 | 0.678 | 0.650 | 0.215 | 0.562 | 1.000 | 0.863 | 0.011 | 0.302 | 0.836 | 0.000 | 0.402 | 0.648 | 0.550 |
| PCA | ✓ | 0.568 | 0.797 | 0.701 | 0.691 | 0.261 | 0.582 | 1.000 | 0.879 | 0.021 | 0.377 | 0.759 | 0.000 | 0.407 | 0.657 | 0.557 |
| TSNE |  | 0.455 | 0.607 | 0.106 | 0.603 | 0.180 | 0.523 | 1.000 | 0.489 | 0.018 | 0.279 | 0.547 | 0.025 | 0.272 | 0.496 | 0.407 |
| TSNE | ✓ | 0.397 | 0.632 | 0.129 | 0.660 | 0.212 | 0.525 | 1.000 | 0.469 | 0.036 | 0.346 | 0.648 | 0.000 | 0.300 | 0.508 | 0.425 |
| UMAP |  | 0.644 | 0.632 | 0.144 | 0.652 | 0.215 | 0.596 | 1.000 | 0.520 | 0.050 | 0.326 | 0.601 | 0.203 | 0.340 | 0.555 | 0.469 |
| UMAP | ✓ | 0.459 | 0.644 | 0.150 | 0.685 | 0.243 | 0.612 | 1.000 | 0.498 | 0.065 | 0.372 | 0.652 | 0.120 | 0.341 | 0.542 | 0.462 |
| Harmony |  | 0.618 | 0.823 | 0.845 | 0.654 | 0.250 | 0.591 | 1.000 | 0.825 | 0.054 | 0.521 | 0.742 | 0.765 | 0.581 | 0.683 | 0.642 |
| Harmony | ✓ | 0.549 | 0.844 | 0.860 | 0.685 | 0.274 | 0.602 | 1.000 | 0.853 | 0.054 | 0.511 | 0.635 | 0.634 | 0.537 | 0.688 | 0.628 |
| Scanorama |  | 0.636 | 0.764 | 0.678 | 0.670 | 0.250 | 0.569 | 1.000 | 0.888 | 0.025 | 0.385 | 0.713 | 0.023 | 0.407 | 0.652 | 0.554 |
| Scanorama | ✓ | 0.562 | 0.783 | 0.697 | 0.704 | 0.333 | 0.594 | 1.000 | 0.835 | 0.030 | 0.437 | 0.578 | 0.128 | 0.402 | 0.668 | 0.561 |
| BBKNN |  | 0.399 | 0.639 | 0.141 | 0.683 | 0.234 | 0.651 | 1.000 | 0.489 | 0.072 | 0.381 | 0.616 | 0.412 | 0.394 | 0.536 | 0.479 |
| BBKNN | ✓ | 0.317 | 0.653 | 0.154 | 0.696 | 0.247 | 0.677 | 1.000 | 0.468 | 0.076 | 0.405 | 0.656 | 0.271 | 0.375 | 0.535 | 0.471 |
| fastMNN | ✓ | 0.503 | 0.050 | 0.008 | 0.043 | 0.005 | 0.439 | 0.926 | 0.780 | 0.046 | 0.044 | 0.055 | 0.330 | 0.251 | 0.282 | 0.270 |
| scVI |  | 0.618 | 0.828 | 0.739 | 0.697 | 0.255 | 0.565 | 1.000 | 0.837 | 0.028 | 0.387 | 0.892 | 0.306 | 0.490 | 0.672 | 0.599 |
| scVI | ✓ | 0.593 | 0.860 | 0.866 | 0.706 | 0.257 | 0.557 | 1.000 | 0.862 | 0.038 | 0.467 | 0.835 | 0.228 | 0.486 | 0.691 | 0.609 |
| scANVI |  | 0.600 | 0.841 | 0.771 | 0.720 | 0.265 | 0.597 | 1.000 | 0.810 | 0.033 | 0.417 | 0.890 | 0.405 | 0.511 | 0.685 | 0.615 |
| scANVI | ✓ | 0.570 | 0.864 | 0.868 | 0.737 | 0.309 | 0.681 | 1.000 | 0.756 | 0.049 | 0.492 | 0.848 | 0.364 | 0.502 | 0.718 | 0.632 |
| scGen | ✓ | 0.694 | 0.939 | 0.966 | 0.763 | 0.312 | 0.687 | 1.000 | 0.746 | 0.058 | 0.634 | 0.838 | 0.525 | 0.560 | 0.766 | 0.683 |
| scPoli | ✓ | 0.411 | 0.828 | 0.680 | 0.728 | 0.317 | 0.767 | 1.000 | 0.461 | 0.070 | 0.618 | 0.668 | 0.449 | 0.453 | 0.676 | 0.587 |
| Author's |  | 0.613 | 0.836 | 0.750 | 0.736 | 0.320 | 0.644 | 1.000 | 0.735 | 0.045 | 0.533 | 0.776 | 0.587 | 0.535 | 0.700 | 0.634 |
| Geneformer |  | 0.644 | 0.753 | 0.541 | 0.552 | 0.196 | 0.520 | 1.000 | 0.857 | 0.032 | 0.410 | 0.780 | 0.419 | 0.500 | 0.601 | 0.560 |
| Islander |  | 0.773 | 1.000 | 1.000 | 0.890 | 0.617 | 0.838 | 1.000 | 0.754 | 0.070 | 0.733 | 0.965 | 0.707 | 0.646 | 0.874 | 0.783 |

**Extended Data Table 5:** Benchmarking cell embeddings, on COVID Cell Atlas.

|  |  | Bio conservation |  |  |  |  |  |  | Batch correction |  |  |  |  | Aggregate score |  |  |
| --- | --- | --- | --- | --- | --- | --- | --- | --- | --- | --- | --- | --- | --- | --- | --- | --- |
| Method | HVG | I-label | L-NMI | L-ARI | K-NMI | K-ARI | S-label | cLISI | S-batch | iLISI | KBET | G-Con | PCR | Batch | Bio | Total |
| PCA |  | 0.495 | 0.698 | 0.669 | 0.563 | 0.226 | 0.520 | 0.996 | 0.921 | 0.122 | 0.215 | 0.751 | 0.000 | 0.402 | 0.595 | 0.518 |
| PCA | ✓ | 0.427 | 0.732 | 0.690 | 0.579 | 0.223 | 0.516 | 0.996 | 0.932 | 0.227 | 0.261 | 0.630 | 0.000 | 0.410 | 0.595 | 0.521 |
| TSNE |  | 0.232 | 0.522 | 0.114 | 0.544 | 0.187 | 0.435 | 0.994 | 0.724 | 0.149 | 0.125 | 0.608 | 0.213 | 0.364 | 0.433 | 0.405 |
| TSNE | ✓ | 0.185 | 0.520 | 0.112 | 0.562 | 0.204 | 0.442 | 0.994 | 0.737 | 0.292 | 0.090 | 0.560 | 0.152 | 0.366 | 0.431 | 0.405 |
| UMAP |  | 0.224 | 0.519 | 0.125 | 0.560 | 0.221 | 0.487 | 0.990 | 0.743 | 0.353 | 0.172 | 0.561 | 0.088 | 0.383 | 0.447 | 0.421 |
| UMAP | ✓ | 0.156 | 0.523 | 0.123 | 0.576 | 0.221 | 0.498 | 0.991 | 0.748 | 0.407 | 0.134 | 0.533 | 0.000 | 0.364 | 0.441 | 0.410 |
| Harmony |  | 0.485 | 0.697 | 0.674 | 0.563 | 0.218 | 0.522 | 0.996 | 0.936 | 0.319 | 0.411 | 0.711 | 0.609 | 0.597 | 0.593 | 0.595 |
| Harmony | ✓ | 0.420 | 0.725 | 0.668 | 0.578 | 0.220 | 0.517 | 0.995 | 0.938 | 0.320 | 0.457 | 0.590 | 0.514 | 0.564 | 0.589 | 0.579 |
| Scanorama |  | 0.443 | 0.706 | 0.669 | 0.565 | 0.226 | 0.517 | 0.995 | 0.932 | 0.270 | 0.257 | 0.659 | 0.345 | 0.493 | 0.589 | 0.550 |
| Scanorama | ✓ | 0.439 | 0.737 | 0.690 | 0.586 | 0.233 | 0.515 | 0.996 | 0.937 | 0.285 | 0.349 | 0.373 | 0.330 | 0.455 | 0.599 | 0.542 |
| BBKNN |  | 0.214 | 0.518 | 0.119 | 0.568 | 0.208 | 0.497 | 0.990 | 0.761 | 0.414 | 0.270 | 0.589 | 0.252 | 0.457 | 0.445 | 0.450 |
| BBKNN | ✓ | 0.128 | 0.533 | 0.132 | 0.581 | 0.216 | 0.503 | 0.992 | 0.758 | 0.442 | 0.188 | 0.546 | 0.052 | 0.397 | 0.441 | 0.423 |
| fastMNN | ✓ | 0.431 | 0.732 | 0.683 | 0.584 | 0.238 | 0.521 | 0.996 | 0.929 | 0.279 | 0.353 | 0.584 | 0.340 | 0.497 | 0.598 | 0.558 |
| scVI |  | 0.501 | 0.703 | 0.657 | 0.588 | 0.261 | 0.519 | 0.997 | 0.933 | 0.228 | 0.259 | 0.788 | 0.404 | 0.522 | 0.604 | 0.571 |
| scVI | ✓ | 0.544 | 0.718 | 0.660 | 0.541 | 0.205 | 0.502 | 0.991 | 0.933 | 0.307 | 0.437 | 0.685 | 0.466 | 0.565 | 0.595 | 0.583 |
| scANVI |  | 0.439 | 0.749 | 0.719 | 0.622 | 0.267 | 0.544 | 0.999 | 0.920 | 0.256 | 0.272 | 0.823 | 0.307 | 0.516 | 0.620 | 0.578 |
| scANVI | ✓ | 0.559 | 0.762 | 0.739 | 0.632 | 0.269 | 0.554 | 0.998 | 0.906 | 0.304 | 0.384 | 0.774 | 0.278 | 0.529 | 0.645 | 0.598 |
| scGen | ✓ | 0.453 | 0.779 | 0.768 | 0.638 | 0.294 | 0.553 | 0.999 | 0.937 | 0.327 | 0.511 | 0.759 | 0.292 | 0.565 | 0.641 | 0.611 |
| scPoli |  | 0.231 | 0.770 | 0.741 | 0.666 | 0.352 | 0.584 | 0.998 | 0.851 | 0.364 | 0.494 | 0.826 | 0.303 | 0.568 | 0.620 | 0.599 |
| scPoli | ✓ | 0.411 | 0.715 | 0.567 | 0.662 | 0.357 | 0.572 | 0.997 | 0.801 | 0.418 | 0.583 | 0.769 | 0.242 | 0.562 | 0.612 | 0.592 |
| Islander |  | 0.693 | 0.999 | 1.000 | 0.863 | 0.603 | 0.713 | 1.000 | 0.934 | 0.420 | 0.653 | 0.967 | 0.693 | 0.733 | 0.839 | 0.797 |

**Extended Data Table 6:** Benchmarking cell embeddings, on Eye Cell Atlas.

|  |  | Bio conservation |  |  |  |  |  |  | Batch correction |  |  |  |  | Aggregate score |  |  |
| --- | --- | --- | --- | --- | --- | --- | --- | --- | --- | --- | --- | --- | --- | --- | --- | --- |
| Method | HVG | I-label | L-NMI | L-ARI | K-NMI | K-ARI | S-label | cLISI | S-batch | iLISI | kBET | G-Con | PCR | Batch | Bio | Total |
| PCA |  | 0.629 | 0.677 | 0.333 | 0.691 | 0.414 | 0.687 | 1.000 | 0.905 | 0.108 | 0.090 | 0.751 | 0.000 | 0.371 | 0.633 | 0.528 |
| PCA | ✓ | 0.637 | 0.742 | 0.550 | 0.701 | 0.415 | 0.705 | 1.000 | 0.903 | 0.175 | 0.207 | 0.662 | 0.000 | 0.390 | 0.678 | 0.563 |
| TSNE |  | 0.553 | 0.529 | 0.086 | 0.481 | 0.177 | 0.521 | 1.000 | 0.770 | 0.105 | 0.015 | 0.357 | 0.000 | 0.250 | 0.478 | 0.387 |
| TSNE | ✓ | 0.563 | 0.541 | 0.088 | 0.512 | 0.203 | 0.551 | 1.000 | 0.777 | 0.164 | 0.031 | 0.538 | 0.000 | 0.302 | 0.494 | 0.417 |
| UMAP |  | 0.569 | 0.538 | 0.087 | 0.583 | 0.309 | 0.594 | 1.000 | 0.785 | 0.121 | 0.031 | 0.572 | 0.000 | 0.302 | 0.526 | 0.436 |
| UMAP | ✓ | 0.587 | 0.550 | 0.095 | 0.625 | 0.389 | 0.707 | 1.000 | 0.773 | 0.180 | 0.097 | 0.603 | 0.000 | 0.331 | 0.565 | 0.471 |
| Harmony |  | 0.644 | 0.769 | 0.649 | 0.701 | 0.393 | 0.721 | 1.000 | 0.900 | 0.360 | 0.227 | 0.802 | 0.855 | 0.629 | 0.697 | 0.670 |
| Harmony | ✓ | 0.641 | 0.855 | 0.967 | 0.773 | 0.546 | 0.718 | 1.000 | 0.900 | 0.389 | 0.273 | 0.643 | 0.626 | 0.566 | 0.786 | 0.698 |
| Scanorama |  | 0.622 | 0.748 | 0.548 | 0.754 | 0.551 | 0.678 | 1.000 | 0.903 | 0.236 | 0.187 | 0.730 | 0.621 | 0.535 | 0.700 | 0.634 |
| Scanorama | ✓ | 0.639 | 0.836 | 0.964 | 0.675 | 0.319 | 0.695 | 1.000 | 0.907 | 0.320 | 0.341 | 0.504 | 0.459 | 0.506 | 0.733 | 0.642 |
| BBKNN |  | 0.617 | 0.545 | 0.094 | 0.544 | 0.238 | 0.706 | 1.000 | 0.762 | 0.275 | 0.079 | 0.628 | 0.571 | 0.463 | 0.535 | 0.506 |
| BBKNN | ✓ | 0.594 | 0.541 | 0.093 | 0.643 | 0.378 | 0.788 | 1.000 | 0.734 | 0.293 | 0.127 | 0.600 | 0.175 | 0.386 | 0.577 | 0.500 |
| fastMNN | ✓ | 0.483 | 0.007 | 0.009 | 0.004 | 0.020 | 0.492 | 0.844 | 0.888 | 0.317 | 0.129 | 0.050 | 0.586 | 0.394 | 0.266 | 0.317 |
| scVI |  | 0.560 | 0.850 | 0.968 | 0.432 | 0.171 | 0.565 | 1.000 | 0.899 | 0.403 | 0.252 | 0.916 | 0.861 | 0.666 | 0.650 | 0.656 |
| scVI | ✓ | 0.620 | 0.845 | 0.965 | 0.619 | 0.237 | 0.526 | 1.000 | 0.916 | 0.431 | 0.307 | 0.786 | 0.594 | 0.607 | 0.688 | 0.655 |
| scANVI |  | 0.597 | 0.862 | 0.972 | 0.729 | 0.528 | 0.624 | 1.000 | 0.890 | 0.400 | 0.237 | 0.918 | 0.866 | 0.662 | 0.759 | 0.720 |
| scANVI | ✓ | 0.685 | 0.775 | 0.591 | 0.776 | 0.433 | 0.650 | 1.000 | 0.886 | 0.459 | 0.382 | 0.630 | 0.725 | 0.616 | 0.701 | 0.667 |
| scGen | ✓ | 0.689 | 0.862 | 0.968 | 0.777 | 0.547 | 0.764 | 1.000 | 0.893 | 0.455 | 0.363 | 0.805 | 0.655 | 0.634 | 0.801 | 0.734 |
| scPoli |  | 0.716 | 0.924 | 0.980 | 0.975 | 0.983 | 0.889 | 1.000 | 0.849 | 0.370 | 0.365 | 0.930 | 0.849 | 0.672 | 0.924 | 0.823 |
| scPoli | ✓ | 0.872 | 0.819 | 0.632 | 0.852 | 0.651 | 0.915 | 1.000 | 0.767 | 0.360 | 0.525 | 0.914 | 0.570 | 0.627 | 0.820 | 0.743 |
| Author's |  | 0.649 | 0.582 | 0.102 | 0.710 | 0.400 | 0.808 | 1.000 | 0.791 | 0.503 | 0.386 | 0.814 | 0.740 | 0.647 | 0.607 | 0.623 |
| Geneformer |  | 0.513 | 0.617 | 0.439 | 0.443 | 0.218 | 0.559 | 1.000 | 0.928 | 0.316 | 0.211 | 0.789 | 0.000 | 0.449 | 0.541 | 0.504 |
| Islander |  | 0.906 | 1.000 | 1.000 | 0.982 | 0.996 | 0.920 | 1.000 | 0.922 | 0.492 | 0.707 | 0.979 | 0.849 | 0.790 | 0.972 | 0.899 |

**Extended Data Table 7:**
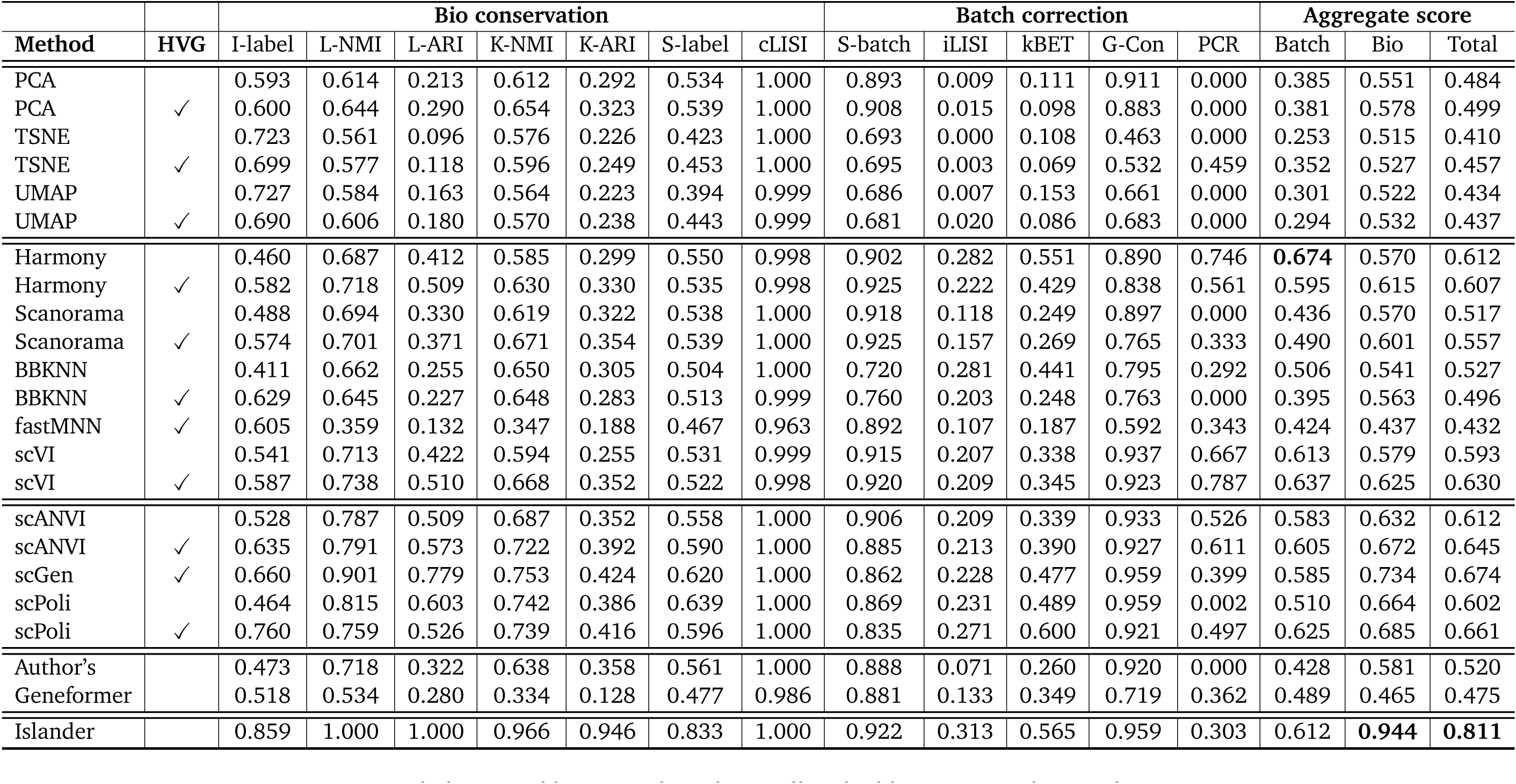
Benchmarking cell embeddings, on Fetal Gut Atlas.

**Extended Data Table 8:**
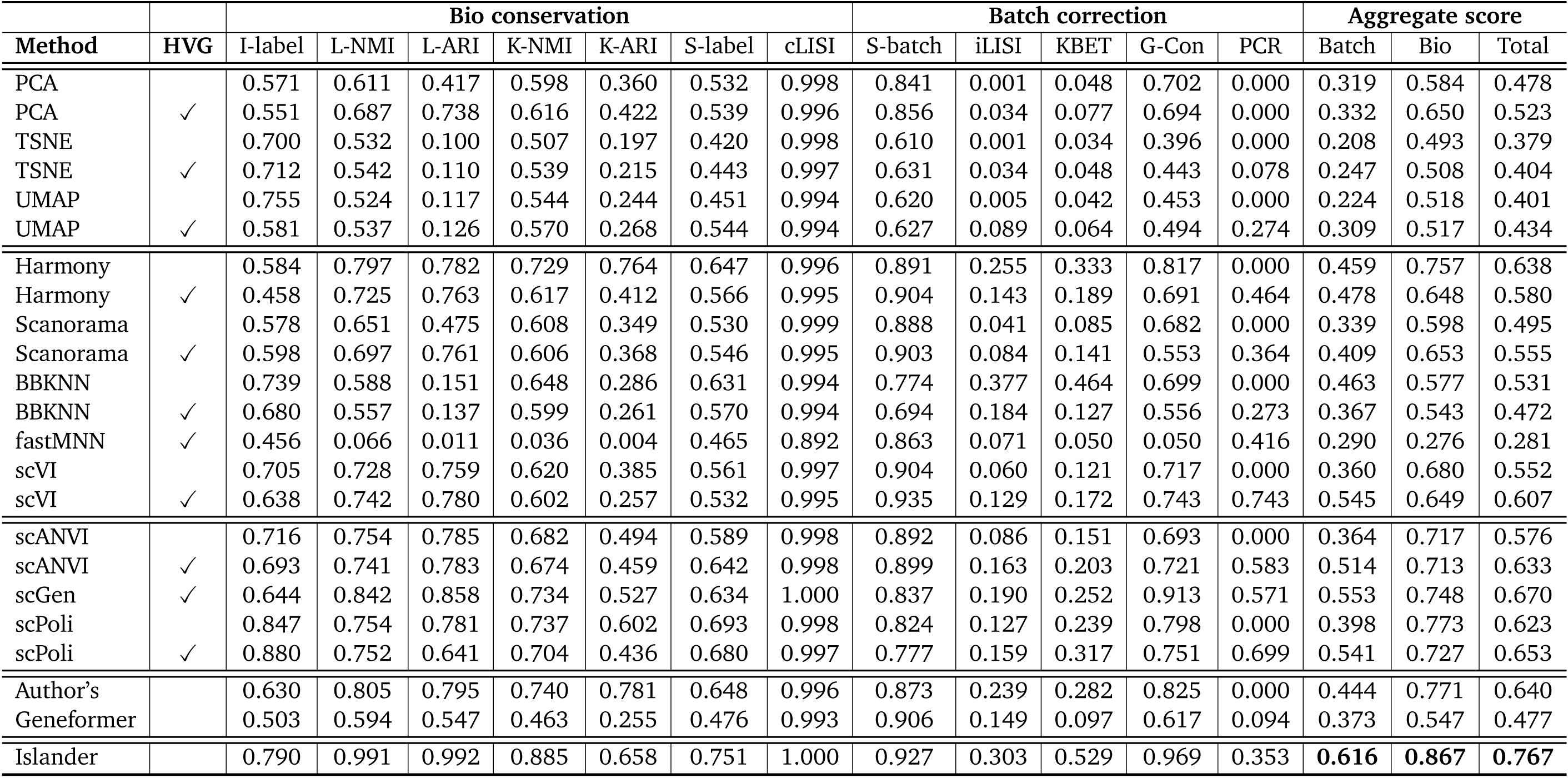
Benchmarking cell embeddings, on Heart Cell Atlas.

**Extended Data Table 9:**
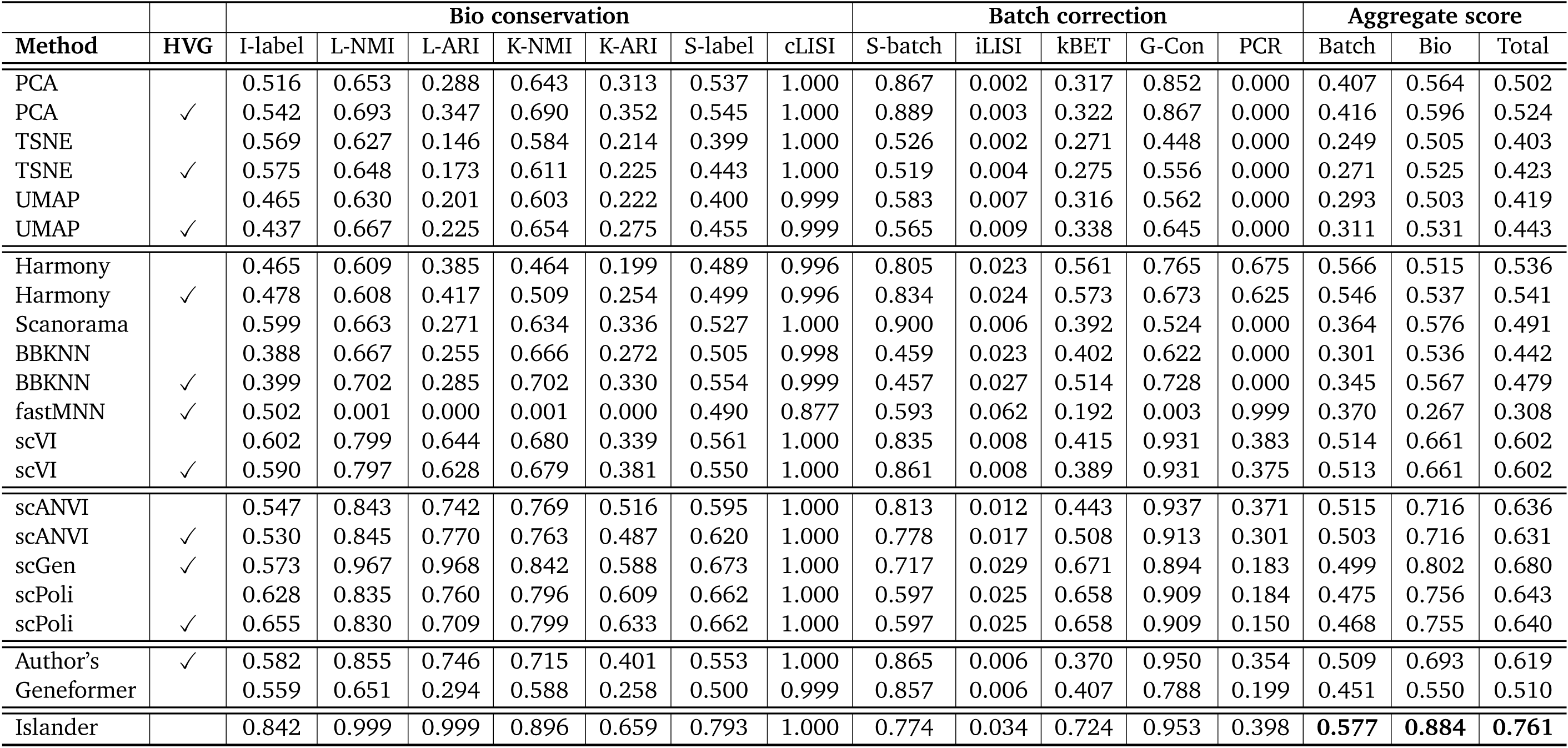
Benchmarking cell embeddings, on Lung Cell Atlas.

**Extended Data Table 10:**
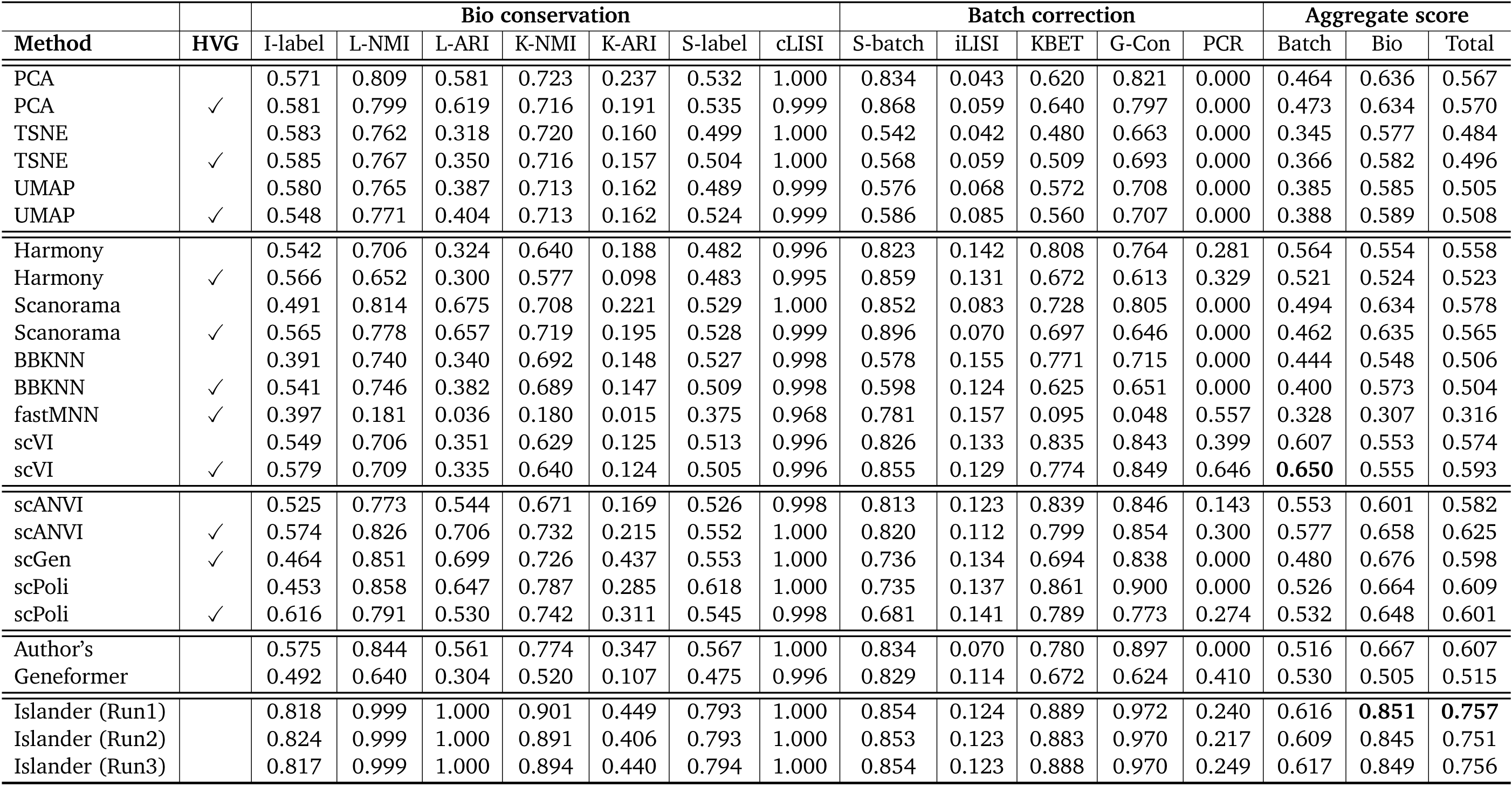
Benchmarking cell embeddings, on (Lung, Fetal, Donor) Cell Atlas.

**Extended Data Table 11:** Benchmarking cell embeddings, on (Lung, Fetal, Organoid) Cell Atlas.

|  |  | Bio conservation |  |  |  |  |  |  | Batch correction |  |  |  |  | Aggregate score |  |  |
| --- | --- | --- | --- | --- | --- | --- | --- | --- | --- | --- | --- | --- | --- | --- | --- | --- |
| Method | HVG | I-label | L-NMI | L-ARI | K-NMI | K-ARI | S-label | cLISI | S-batch | iLISI | kBET | G-Con | PCR | Batch | Bio | Total |
| PCA |  | 0.466 | 0.441 | 0.264 | 0.399 | 0.126 | 0.512 | 0.992 | 0.791 | 0.000 | 0.486 | 0.728 | 0.000 | 0.401 | 0.457 | 0.435 |
| PCA | ✓ | 0.439 | 0.451 | 0.235 | 0.402 | 0.125 | 0.506 | 0.992 | 0.821 | 0.000 | 0.521 | 0.738 | 0.000 | 0.416 | 0.450 | 0.436 |
| TSNE |  | 0.281 | 0.378 | 0.061 | 0.381 | 0.093 | 0.430 | 0.991 | 0.522 | 0.000 | 0.341 | 0.513 | 0.000 | 0.275 | 0.374 | 0.334 |
| TSNE | ✓ | 0.313 | 0.378 | 0.068 | 0.391 | 0.095 | 0.420 | 0.990 | 0.564 | 0.000 | 0.332 | 0.558 | 0.000 | 0.291 | 0.379 | 0.344 |
| UMAP |  | 0.294 | 0.396 | 0.105 | 0.396 | 0.102 | 0.429 | 0.985 | 0.546 | 0.000 | 0.402 | 0.587 | 0.000 | 0.307 | 0.387 | 0.355 |
| UMAP | ✓ | 0.382 | 0.393 | 0.097 | 0.398 | 0.111 | 0.468 | 0.986 | 0.578 | 0.000 | 0.491 | 0.647 | 0.000 | 0.343 | 0.405 | 0.380 |
| Harmony |  | 0.546 | 0.324 | 0.177 | 0.267 | 0.084 | 0.465 | 0.982 | 0.793 | 0.034 | 0.607 | 0.466 | 0.276 | 0.435 | 0.406 | 0.418 |
| Harmony | ✓ | 0.484 | 0.336 | 0.148 | 0.215 | 0.045 | 0.428 | 0.982 | 0.870 | 0.017 | 0.621 | 0.608 | 0.712 | 0.565 | 0.377 | 0.452 |
| Scanorama |  | 0.532 | 0.441 | 0.298 | 0.359 | 0.105 | 0.484 | 0.991 | 0.837 | 0.000 | 0.477 | 0.698 | 0.000 | 0.403 | 0.459 | 0.436 |
| Scanorama | ✓ | 0.442 | 0.430 | 0.249 | 0.386 | 0.119 | 0.488 | 0.991 | 0.824 | 0.000 | 0.548 | 0.595 | 0.081 | 0.410 | 0.444 | 0.430 |
| BBKNN |  | 0.434 | 0.374 | 0.094 | 0.363 | 0.091 | 0.438 | 0.981 | 0.593 | 0.036 | 0.462 | 0.632 | 0.000 | 0.345 | 0.396 | 0.376 |
| BBKNN | ✓ | 0.396 | 0.346 | 0.079 | 0.357 | 0.090 | 0.367 | 0.980 | 0.626 | 0.017 | 0.323 | 0.336 | 0.000 | 0.260 | 0.374 | 0.328 |
| fastMNN | ✓ | 0.440 | 0.254 | 0.111 | 0.161 | 0.035 | 0.444 | 0.955 | 0.846 | 0.006 | 0.106 | 0.131 | 0.463 | 0.310 | 0.343 | 0.330 |
| scVI |  | 0.485 | 0.344 | 0.251 | 0.164 | 0.032 | 0.457 | 0.977 | 0.870 | 0.024 | 0.581 | 0.706 | 0.821 | 0.600 | 0.387 | 0.473 |
| scVI | ✓ | 0.517 | 0.283 | 0.136 | 0.183 | 0.040 | 0.452 | 0.970 | 0.880 | 0.031 | 0.557 | 0.722 | 0.934 | 0.625 | 0.369 | 0.471 |
| scANVI |  | 0.473 | 0.471 | 0.360 | 0.320 | 0.095 | 0.458 | 0.992 | 0.854 | 0.018 | 0.576 | 0.708 | 0.622 | 0.555 | 0.453 | 0.494 |
| scANVI | ✓ | 0.508 | 0.455 | 0.347 | 0.366 | 0.111 | 0.453 | 0.990 | 0.832 | 0.013 | 0.588 | 0.751 | 0.737 | 0.584 | 0.462 | 0.511 |
| scGen | ✓ | 0.572 | 0.908 | 0.890 | 0.649 | 0.243 | 0.642 | 1.000 | 0.714 | 0.019 | 0.663 | 0.895 | 0.364 | 0.531 | 0.701 | 0.633 |
| scPoli |  | 0.631 | 0.477 | 0.301 | 0.415 | 0.115 | 0.485 | 0.993 | 0.734 | 0.010 | 0.581 | 0.845 | 0.000 | 0.434 | 0.488 | 0.466 |
| scPoli | ✓ | 0.479 | 0.484 | 0.298 | 0.427 | 0.149 | 0.511 | 0.989 | 0.666 | 0.010 | 0.610 | 0.759 | 0.113 | 0.432 | 0.477 | 0.459 |
| Author's |  | 0.647 | 0.480 | 0.272 | 0.442 | 0.192 | 0.555 | 0.995 | 0.796 | 0.002 | 0.527 | 0.769 | 0.000 | 0.419 | 0.512 | 0.475 |
| Geneformer |  | 0.436 | 0.337 | 0.208 | 0.242 | 0.074 | 0.477 | 0.980 | 0.828 | 0.012 | 0.466 | 0.457 | 0.598 | 0.472 | 0.394 | 0.425 |
| Islander |  | 0.785 | 0.955 | 0.890 | 0.736 | 0.313 | 0.752 | 1.000 | 0.844 | 0.027 | 0.687 | 0.978 | 0.000 | 0.507 | 0.776 | 0.668 |

**Extended Data Table 12:** Benchmarking cell embeddings, on Pancreas Cell Atlas.

|  |  | Bio conservation |  |  |  |  |  |  | Batch correction |  |  |  |  | Aggregate score |  |  |
| --- | --- | --- | --- | --- | --- | --- | --- | --- | --- | --- | --- | --- | --- | --- | --- | --- |
| Method | HVG | I-label | L-NMI | L-ARI | K-NMI | K-ARI | S-label | cLISI | S-batch | iLISI | kBET | G-Con | PCR | Batch | Bio | Total |
| PCA |  | 0.496 | 0.594 | 0.214 | 0.459 | 0.313 | 0.499 | 1.000 | 0.770 | 0.000 | 0.057 | 0.553 | 0.000 | 0.276 | 0.511 | 0.417 |
| PCA | ✓ | 0.634 | 0.685 | 0.421 | 0.620 | 0.408 | 0.572 | 1.000 | 0.849 | 0.002 | 0.206 | 0.656 | 0.000 | 0.343 | 0.620 | 0.509 |
| TSNE |  | 0.363 | 0.573 | 0.161 | 0.438 | 0.203 | 0.417 | 1.000 | 0.482 | 0.000 | 0.030 | 0.424 | 0.000 | 0.187 | 0.451 | 0.345 |
| TSNE | ✓ | 0.954 | 0.634 | 0.236 | 0.589 | 0.340 | 0.451 | 1.000 | 0.546 | 0.000 | 0.060 | 0.550 | 0.000 | 0.231 | 0.601 | 0.453 |
| UMAP |  | 0.372 | 0.580 | 0.211 | 0.387 | 0.167 | 0.386 | 1.000 | 0.511 | 0.000 | 0.036 | 0.555 | 0.000 | 0.220 | 0.443 | 0.354 |
| UMAP | ✓ | 0.932 | 0.630 | 0.240 | 0.589 | 0.343 | 0.496 | 1.000 | 0.570 | 0.000 | 0.114 | 0.587 | 0.000 | 0.254 | 0.604 | 0.464 |
| Harmony |  | 0.553 | 0.869 | 0.908 | 0.623 | 0.426 | 0.570 | 1.000 | 0.885 | 0.198 | 0.421 | 0.853 | 0.918 | 0.655 | 0.707 | 0.686 |
| Harmony | ✓ | 0.586 | 0.904 | 0.945 | 0.792 | 0.647 | 0.619 | 1.000 | 0.868 | 0.220 | 0.538 | 0.765 | 0.790 | 0.636 | 0.785 | 0.725 |
| Scanorama |  | 0.555 | 0.754 | 0.564 | 0.614 | 0.392 | 0.556 | 1.000 | 0.905 | 0.037 | 0.278 | 0.863 | 0.674 | 0.551 | 0.633 | 0.601 |
| Scanorama | ✓ | 0.609 | 0.878 | 0.916 | 0.762 | 0.598 | 0.597 | 1.000 | 0.934 | 0.144 | 0.384 | 0.694 | 0.673 | 0.566 | 0.766 | 0.686 |
| BBKNN |  | 0.473 | 0.623 | 0.271 | 0.626 | 0.367 | 0.547 | 1.000 | 0.660 | 0.023 | 0.190 | 0.676 | 0.671 | 0.444 | 0.558 | 0.512 |
| BBKNN | ✓ | 0.923 | 0.681 | 0.319 | 0.723 | 0.461 | 0.695 | 1.000 | 0.669 | 0.062 | 0.217 | 0.850 | 0.628 | 0.485 | 0.686 | 0.606 |
| fastMNN | ✓ | 0.429 | 0.099 | 0.023 | 0.035 | 0.015 | 0.475 | 0.841 | 0.792 | 0.190 | 0.004 | 0.150 | 0.853 | 0.398 | 0.274 | 0.323 |
| scVI |  | 0.700 | 0.880 | 0.916 | 0.635 | 0.442 | 0.558 | 0.999 | 0.851 | 0.209 | 0.424 | 0.913 | 0.964 | 0.672 | 0.733 | 0.709 |
| scVI | ✓ | 0.664 | 0.920 | 0.953 | 0.743 | 0.451 | 0.573 | 1.000 | 0.881 | 0.277 | 0.527 | 0.910 | 0.856 | 0.690 | 0.757 | 0.731 |
| scANVI |  | 0.720 | 0.917 | 0.949 | 0.732 | 0.712 | 0.572 | 1.000 | 0.844 | 0.207 | 0.411 | 0.923 | 0.955 | 0.668 | 0.800 | 0.747 |
| scANVI | ✓ | 0.664 | 0.920 | 0.953 | 0.743 | 0.451 | 0.573 | 1.000 | 0.881 | 0.277 | 0.527 | 0.910 | 0.856 | 0.690 | 0.757 | 0.731 |
| scGen | ✓ | 0.813 | 0.960 | 0.981 | 0.833 | 0.638 | 0.700 | 1.000 | 0.777 | 0.286 | 0.637 | 0.938 | 0.754 | 0.678 | 0.847 | 0.779 |
| scPoli |  | 0.786 | 0.919 | 0.949 | 0.783 | 0.541 | 0.701 | 1.000 | 0.822 | 0.192 | 0.485 | 0.931 | 0.871 | 0.660 | 0.811 | 0.751 |
| scPoli | ✓ | 0.682 | 0.912 | 0.945 | 0.829 | 0.733 | 0.760 | 1.000 | 0.758 | 0.344 | 0.677 | 0.939 | 0.757 | 0.695 | 0.837 | 0.780 |
| Islander |  | 0.893 | 0.999 | 0.999 | 0.980 | 0.984 | 0.914 | 1.000 | 0.873 | 0.254 | 0.558 | 0.977 | 0.897 | 0.712 | 0.967 | 0.865 |

**Extended Data Table 13:**
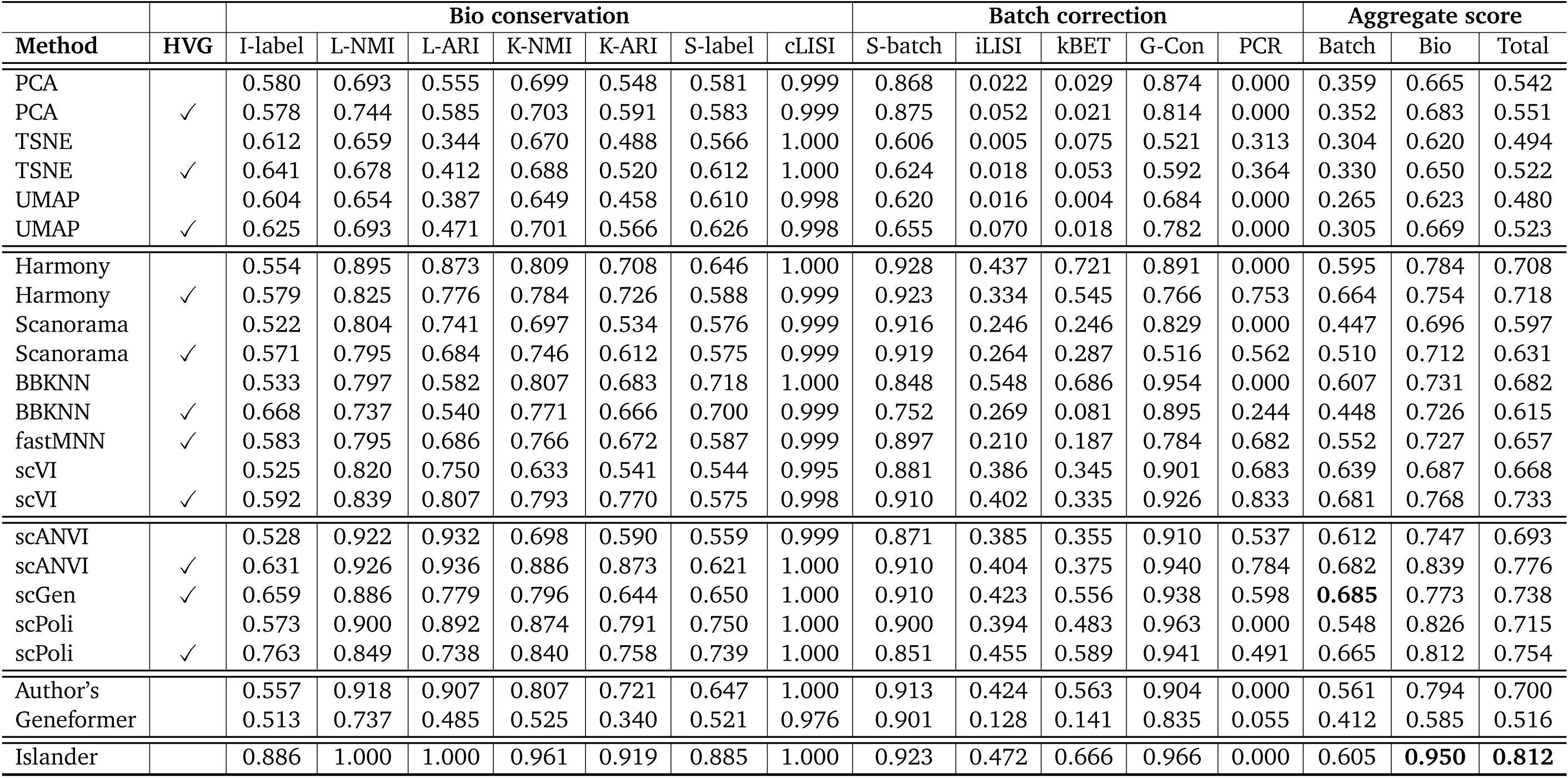
Benchmarking cell embeddings, on Skin Cell Atlas.

**Extended Data Table 14:**
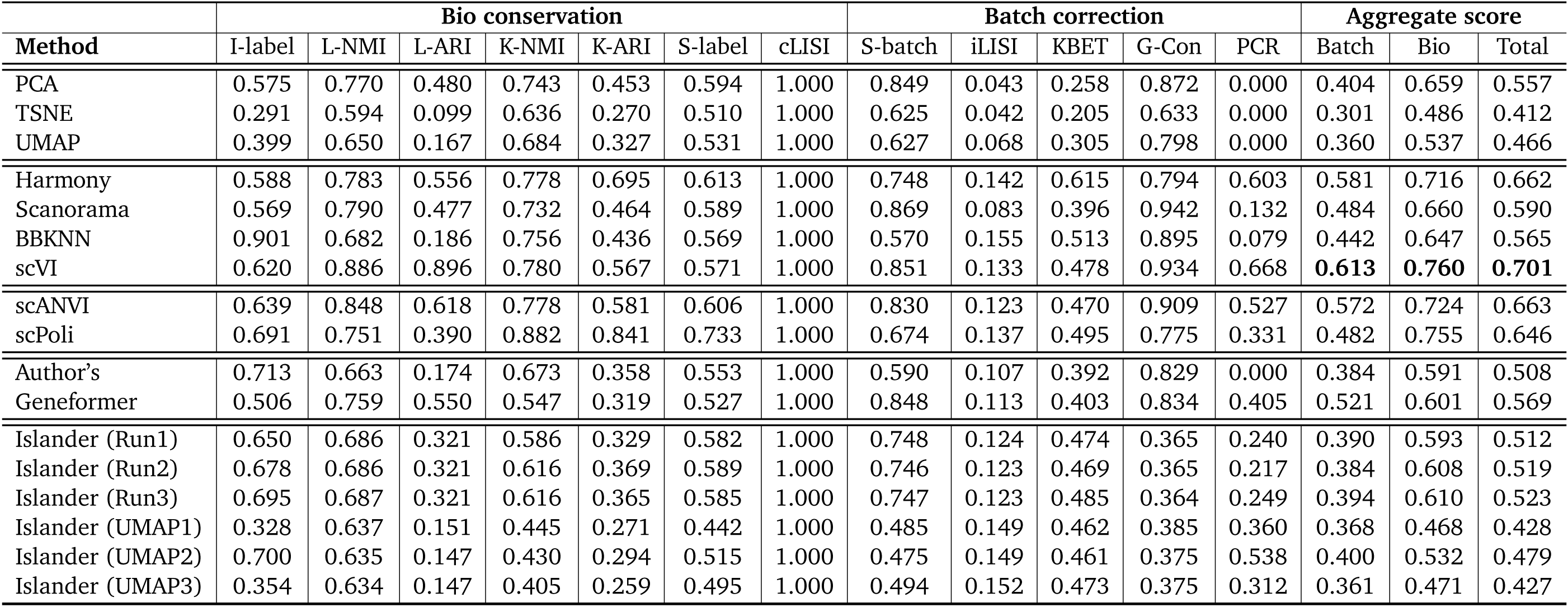
Benchmarking cell embeddings, on (Lung, Fetal, Donor) Cell Atlas, using broad cell types.

**Extended Data Table 15:** Benchmarking cell embeddings, using scGraph. “F”, “D” and “O” represents fetal, donor and organoid, respectively. “nan” means the embeddings are not available, due to memory limitations (>500G in RAM) or unavailability of raw counts (Geneformer). We <u>underline</u> the worst methods among integration methods, Geneformer and Islander , for each atlas.

| Method | HVG | Brain | Breast | COVID | Eye | Gut (F) | Heart | Lung (F,D) | Lung (F,O) | Lung | Pancreas | Skin |
| --- | --- | --- | --- | --- | --- | --- | --- | --- | --- | --- | --- | --- |
| PCA |  | 0.932 | 0.885 | 0.808 | 0.941 | 0.968 | 0.746 | 0.769 | 0.707 | 0.827 | 0.909 | 0.914 |
| PCA | ✓ | 0.949 | 0.938 | 0.962 | 0.979 | 0.979 | 0.713 | 0.769 | 0.682 | 0.910 | 0.952 | 0.953 |
| TSNE |  | 0.620 | 0.504 | 0.599 | 0.745 | 0.724 | 0.634 | 0.448 | 0.373 | 0.481 | 0.730 | 0.708 |
| TSNE | ✓ | 0.633 | 0.632 | 0.616 | 0.694 | 0.699 | 0.657 | 0.448 | 0.426 | 0.655 | 0.608 | 0.718 |
| UMAP |  | 0.580 | 0.592 | 0.649 | 0.695 | 0.652 | 0.532 | 0.563 | 0.462 | 0.635 | 0.621 | 0.637 |
| UMAP | ✓ | 0.684 | 0.617 | 0.726 | 0.762 | 0.634 | 0.618 | 0.563 | 0.354 | 0.652 | 0.624 | 0.703 |
| Harmony |  | 0.924 | 0.908 | 0.837 | 0.946 | 0.636 | 0.835 | 0.421 | 0.491 | 0.746 | 0.928 | 0.721 |
| Harmony | ✓ | 0.941 | 0.957 | 0.971 | 0.981 | 0.978 | 0.766 | 0.731 | 0.703 | 0.890 | 0.954 | 0.973 |
| Scanorama |  | 0.929 | 0.872 | 0.813 | 0.949 | 0.934 | 0.735 | 0.669 | 0.689 | 0.738 | 0.894 | 0.922 |
| Scanorama | ✓ | 0.915 | 0.916 | 0.930 | 0.924 | 0.946 | 0.727 | 0.669 | 0.718 | 0.774 | 0.913 | 0.936 |
| BBKNN |  | 0.544 | 0.600 | 0.772 | 0.674 | 0.645 | 0.677 | 0.462 | 0.453 | 0.654 | 0.667 | 0.733 |
| BBKNN | ✓ | 0.562 | 0.601 | 0.765 | 0.618 | 0.739 | 0.608 | 0.613 | 0.588 | 0.669 | 0.583 | 0.768 |
| scVI |  | 0.820 | 0.731 | 0.622 | 0.782 | 0.726 | 0.718 | 0.440 | 0.678 | 0.632 | 0.633 | 0.677 |
| scVI | ✓ | 0.863 | 0.732 | 0.704 | 0.795 | 0.710 | 0.757 | 0.438 | 0.668 | 0.683 | 0.643 | 0.696 |
| scANVI |  | 0.857 | 0.774 | 0.773 | 0.818 | 0.798 | 0.818 | 0.518 | 0.708 | 0.712 | 0.634 | 0.651 |
| scANVI | ✓ | 0.927 | 0.848 | 0.848 | 0.820 | 0.820 | 0.888 | 0.508 | 0.695 | 0.829 | 0.696 | 0.761 |
| scGen | ✓ | 0.949 | 0.880 | 0.868 | 0.954 | 0.946 | 0.688 | 0.694 | 0.607 | 0.872 | 0.901 | 0.899 |
| scPoli |  | nan | nan | 0.629 | 0.824 | 0.857 | 0.829 | 0.732 | 0.645 | 0.826 | 0.836 | 0.690 |
| scPoli | ✓ | 0.869 | 0.818 | 0.741 | 0.933 | 0.849 | 0.858 | 0.729 | 0.647 | 0.826 | 0.733 | 0.887 |
| Author's |  | 0.644 | 0.884 | nan | 0.593 | 0.676 | 0.829 | 0.379 | 0.337 | 0.634 | nan | 0.706 |
| Geneformer |  | 0.799 | 0.631 | nan | 0.693 | 0.821 | 0.693 | 0.597 | 0.625 | 0.668 | nan | 0.800 |
| Islander |  | 0.549 | 0.464 | 0.480 | 0.753 | 0.633 | 0.577 | 0.256 | 0.697 | 0.513 | 0.676 | 0.654 |

### 3. Visualization

**Extended Data Figure 1:**
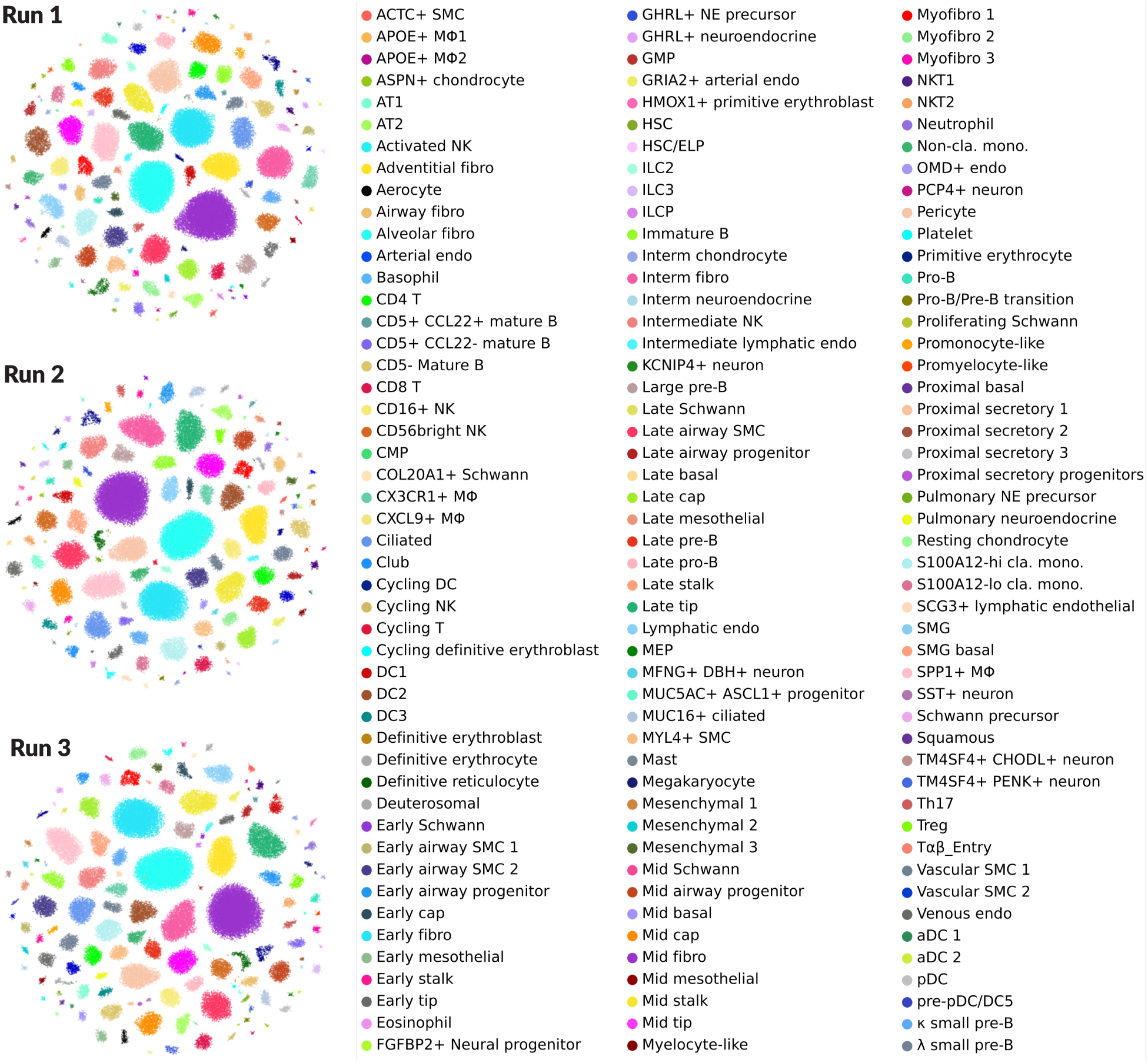
Drifting Cell Islands, different runs of Islander on fetal lung atlas.

**Extended Data Figure 2:**
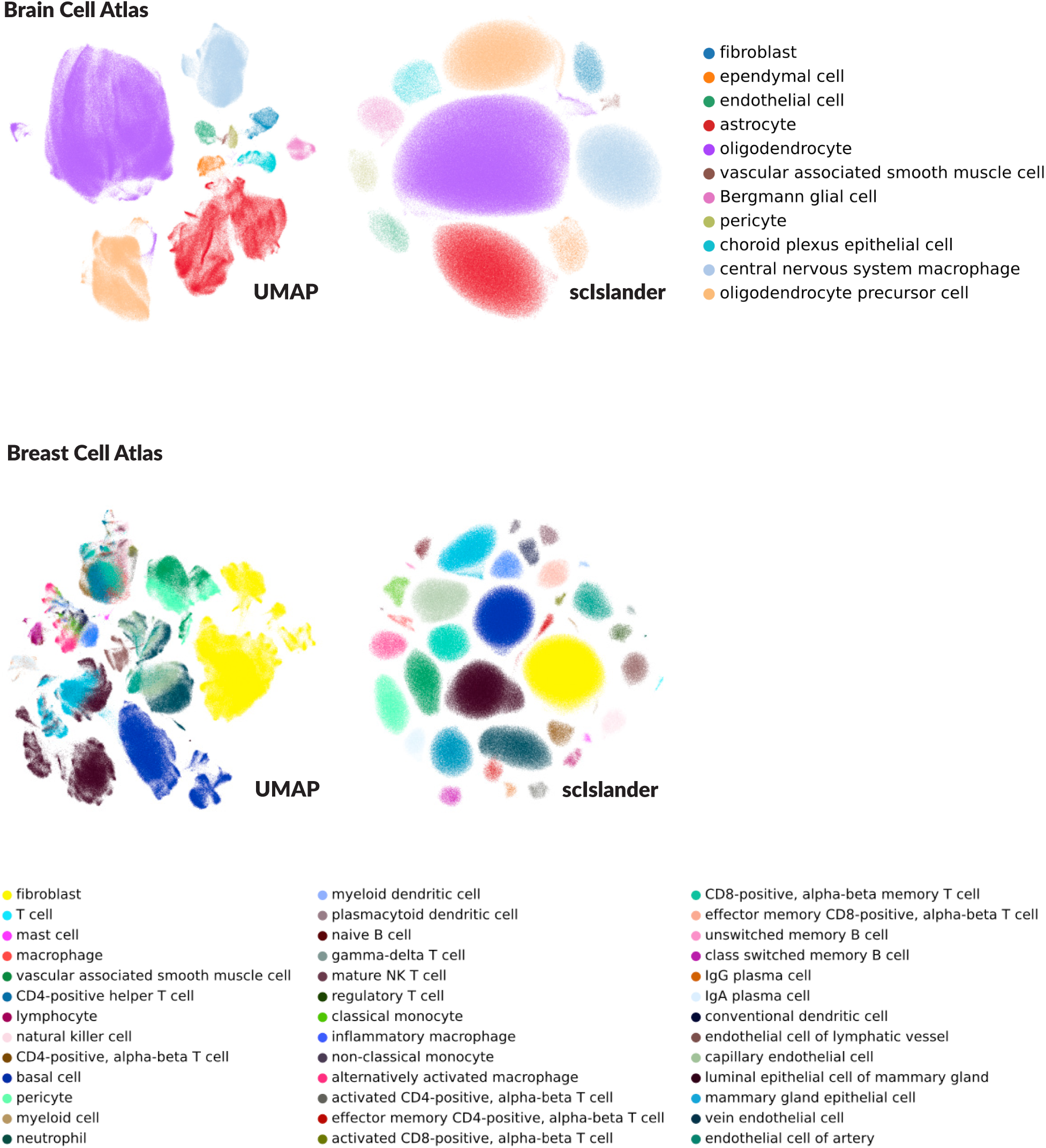
Drifting Cell Islands, Part I.

**Extended Data Figure 3:**
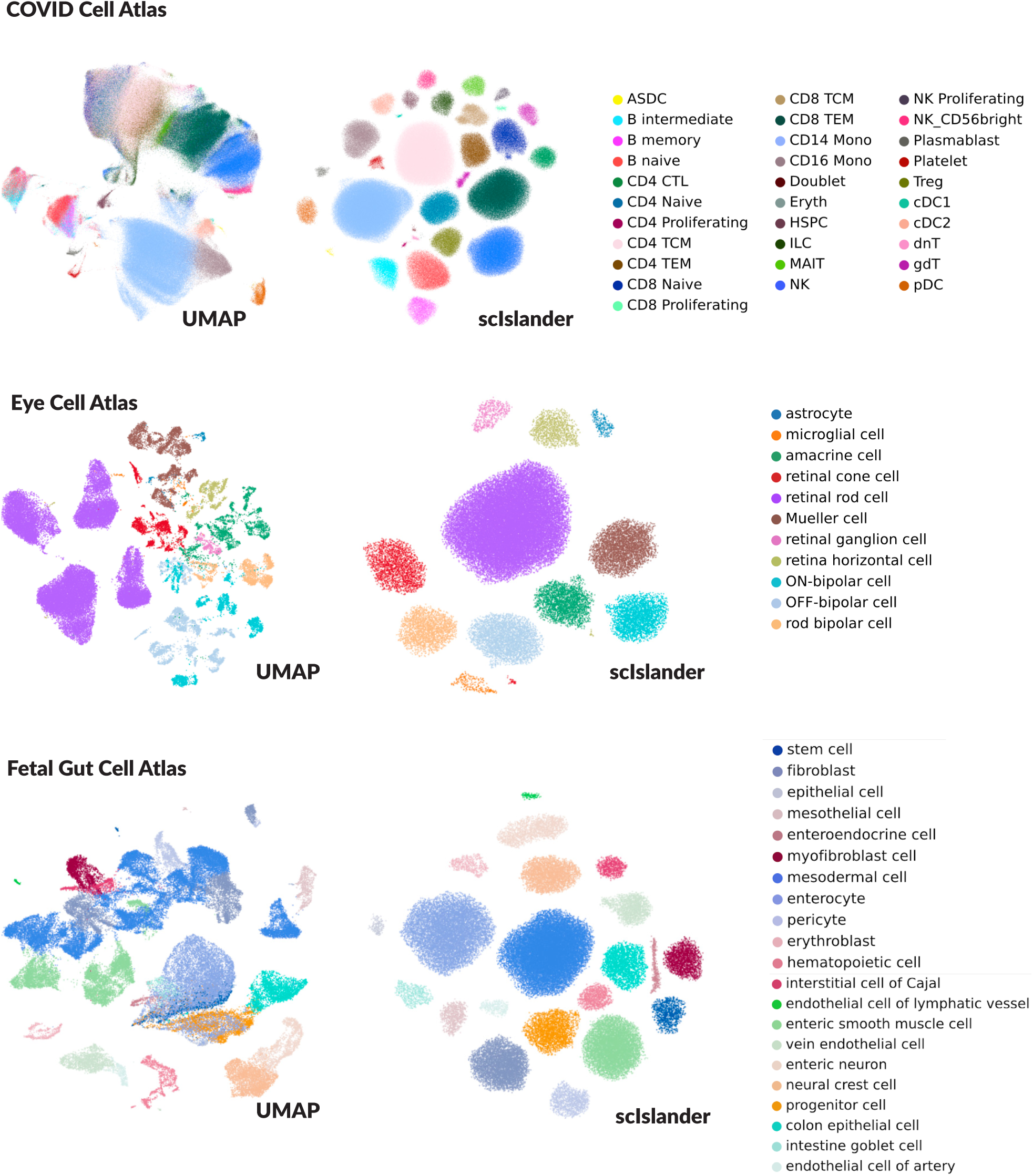
Drifting Cell Islands, Part II.

**Extended Data Figure 4:**
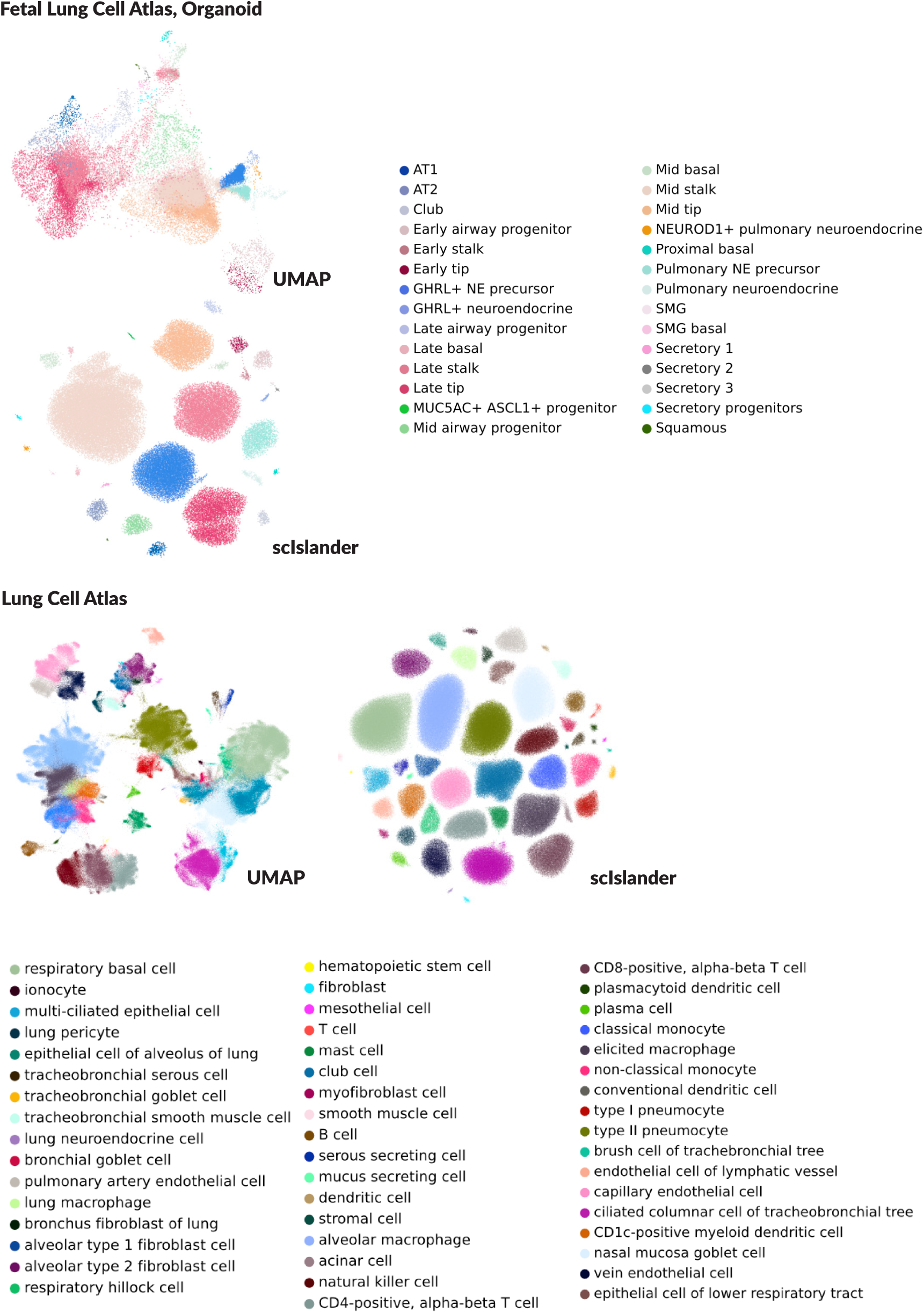
Drifting Cell Islands, Part III.

**Extended Data Figure 5:**
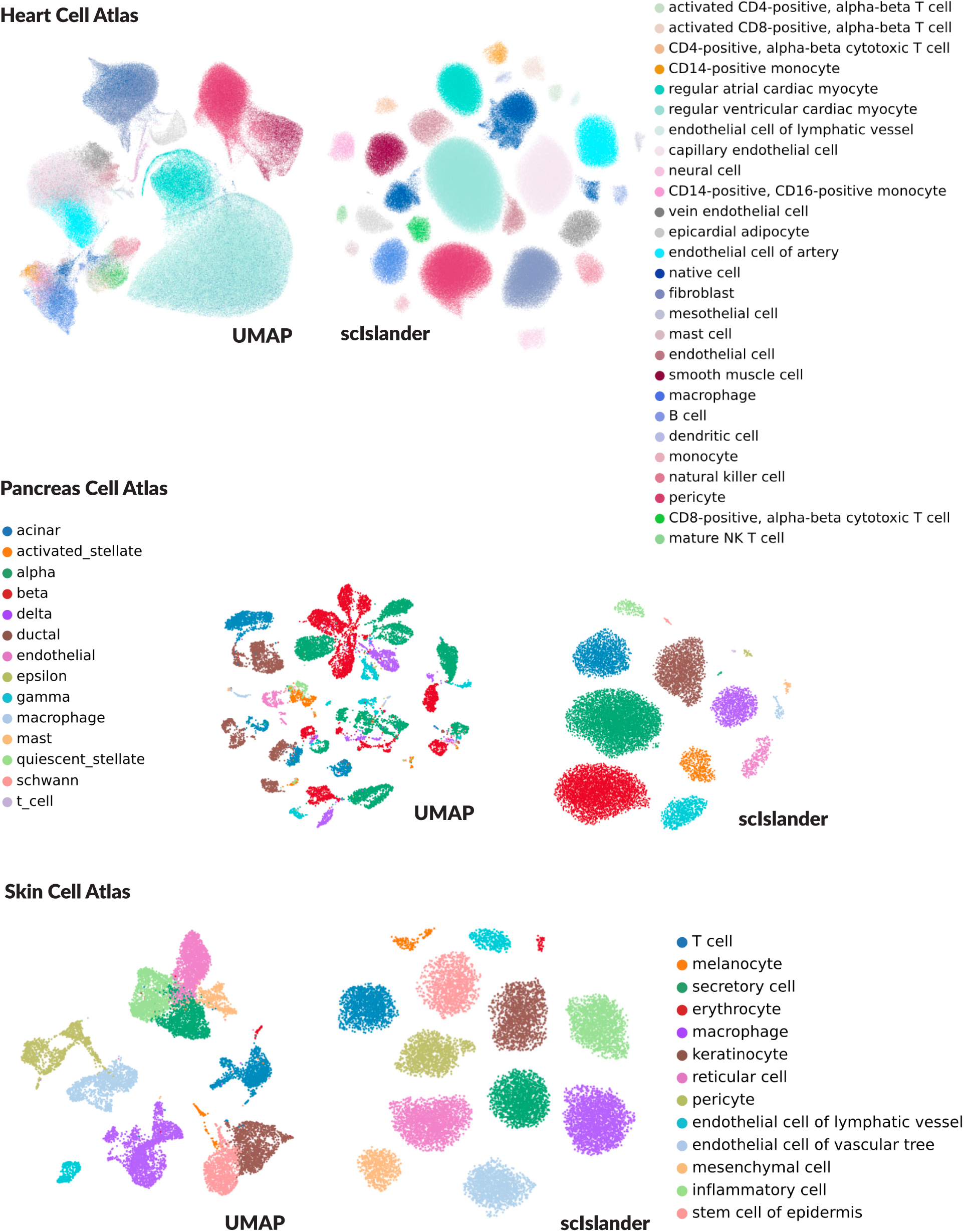
Drifting Cell Islands, Part IV.

